# Sex differences in allometry for mouse phenotypic traits indicate that females are not scaled males

**DOI:** 10.1101/2022.03.29.486193

**Authors:** Laura A. B. Wilson, Susanne R. K. Zajitschek, Malgorzata Lagisz, Jeremy Mason, Hamed Haselimashhadi, Shinichi Nakagawa

**Affiliations:** Evolution & Ecology Research Centre, UNSW Data Science Hub, and School of Biological, Earth and Environmental Sciences, University of New South Wales, Sydney, Australia; School of Archaeology and Anthropology, The Australian National University, Canberra, Australia; School of Biological and Environmental Sciences, Liverpool John Moores University, Liverpool, UK; Melio Healthcare Ltd., City Tower, 40 Basinghall St., EC2V 5DE, London, UK; European Molecular Biology Laboratory, European Bioinformatics Institute (EMBL-EBI), Wellcome Genome Campus, Hinxton, Cambridgeshire, CB10 1SD, UK

**Keywords:** biomarker, sex differences, static allometry, animal model, drug reactions

## Abstract

Sex differences in the lifetime risk and expression of disease are well-known. Preclinical research targeted at improving treatment, increasing health span, and reducing the financial burden of health care, has mostly been conducted on male animals and cells. The extent to which sex differences in phenotypic traits are explained by sex differences in body weight remains unclear. We quantify sex differences in the allometric relationship between trait value and body weight for 375 phenotypic traits in male and female mice, recorded in >2.1 million measurements from the International Mouse Phenotyping Consortium. We find sex differences in allometric parameters (slope, intercept, residual SD) are common (76% traits). Body weight differences do not explain all sex differences in trait values but scaling by weight may be useful for some traits. Our results support a trait-specific patterning of sex differences in phenotypic traits, promoting case-specific approaches to drug dosage scaled by body weight.

## Introduction

A historic use of male animals in preclinical research and male participants in clinical trials has resulted in a significant bias in healthcare systems around the world^1^. The knowledge available on many diseases, their manifestation, time course and the efficacy of treatment options, is highly skewed in favour of males. The need to reach parity of the sexes in biomedical research and to conduct sex-specific analysis of research results has been widely acknowledged^2-4,5,6^. Efforts to address this issue resulted in legislative changes around clinical research, requiring female participants in government-funded clinical trials (e.g.,^7,8,9^). Modest improvement to rebalancing representation of the sexes in clinical trials^10-12^ has been bolstered by recent revisions to government guidelines in the US for preclinical research, requiring biological sex to be included as a study variable^13^.

Preclinical research on mice, one of the most important animal models for investigating human disease^14,15^, is fundamental for informing clinical research. These data illuminate clinically relevant pharmacological processes and enable the testing of treatment effects that would present ethical and safety issues in humans^15^. With the growing recognition of the importance of sex in biomedicine, a sharper focus on the topic in mouse data has revealed that some of the initial assumptions and concerns surrounding use of female animals in preclinical research, such as their propensity for greater variation associated with the oestrous cycle, lack empirical support^2,16-18^.

Building on empirical studies that have sought to establish the nature of sex differences in biomedicine and to clarify the assumptions surrounding preclinical research data collected on males and generalized to females^2,18-20^, we here use an allometric framework and large mouse phenotype data set to tackle the unresolved issue of whether sex differences in traits may be explained by sex differences in body weight. The extent to which body weight may eliminate the statistical significance of sex as an independent variable remains unclear^22^, yet is material to debate surrounding how reductions in health disparities may be effectively targeted. Data on allometric scaling relate to one of the most salient aspects of sex differences, those concerning adverse drug reactions (ADRs). Compared to men, women experience ADRs almost twice as often^23^, with the same therapeutic dose often being prescribed to both sexes^24^. Defining the allometric relationship between phenotypic trait and body weight for males and females is required to better understand whether this relationship is upheld across diverse traits and whether most observed differences are due to scaling. This will help inform whether the majority of sex-specific ADRs might be resolved by implementing weight-adjusted doses and, if not, whether the use of weight-adjusted dosing may be helpful in some circumstances at reducing sex-specific ADRs.

Studies have established that the nature of disease experience and benefits of treatment differ between men and women^28-33^, and that for many systems the observed sex differences in traits relate to differences in underlying processes rather than sex differences in body weight or size. These differences manifest in major pillars of healthcare, impacting cost associated with care and its quality^34^. For example, the broadly divergent behaviour of male (anti-inflammatory) and female (pro-inflammatory) immune systems translates to antibody response variability, with some vaccines resulting in a stronger immune response in males compared to females^37-40^. Further, sex differences in metabolism underlie more pronounced expression of metabolic syndrome disease (e.g., obesity) phenotypes in males^41^, which have been linked to fundamental regulatory differences in metabolic homeostasis, impacting energy partitioning and storage. Pathophysiological differences between the sexes have also been recognised in cardiology, such that diagnostic data extracted from coronary angiograms are interpreted in a sex-specific manner^42^. In the context of drug treatment, sex differences in pharmacokinetics, related to absorption rates (e.g., gastric enzymes^43^), distribution mechanisms (e.g., plasma binding capacity^44^), and metabolism (e.g., renal elimination capacity) result in males having lower free drug concentration and higher drug clearance compared to females^44^. Likewise, sex differences in major renal functions (e.g., glomerular filtration rate^45^) mean that the effects of a drug on the body are also different for men and women, which, together with pharmacokinetic parameters, translates to differences in drug efficacy and toxicity^23^. Extending across and beyond many of these systems in which sex differences have been identified, we here provide empirical data on static allometry across phenotypic traits that represent preclinical parameters (e.g., immunology, metabolism, morphology) in a disease model animal (mouse). We aim to clarify if, and the extent to which, trait values for males may be scaled to match those of females.

We adopt the framework of static allometry, the measurement of trait covariation among individuals of different size at the same developmental stage, following Huxley^46,47^ who proposed an equation to model simple allometry. This equation expresses the growth of two traits, *x* and *y*, when regulated by a common growth parameter: *y* = *axb*, or equivalently log *y* = log(*a*) + *b* log (*y*), where the ratios between the components of the growth rates of y and x correspond to intercept log(*a*) and a slope b^48^. We quantify the relationship between phenotypic trait and body weight in males and females, statistically evaluating scenarios that describe the magnitude and patterning of sex differences across 375 traits in over 2.1 million mice from the International Mouse Phenotyping Consortium^49^ (IMPC, www.mousephenotype.org). We discuss these data considering the discourse on the generalization of male data in preclinical research^50^, as well as their evolutionary implications, leveraging a large, wildtype dataset to illuminate microevolutionary trends in static allometry. Consideration of the evolutionary context surrounding sex differences may augment understanding of how disease state phenotypes emerge or persist in a population^51,52^.

## Results

### Data characteristics

Following initial data cleaning and filtering procedures, the dataset comprised 375 phenotypic traits with a median sample size of 1,506 mice per trait (n = 2,105,419). Representation of males and females was highly similar across most phenotypic traits, with fewer than 15% of traits (53/375) displaying greater than 10% difference in sample size between males and females. The traits were collated into nine functional groupings following Zajitschek et al. (2020) (see Methods): behaviour (85 traits, n = 440,491), eye (39 traits, n = 21,695), hearing (21 traits, n = 273,715), heart (31 traits, n = 233,772), hematology (25 traits, n = 293,312), immunology (111 traits, n = 108,471), metabolism (8 traits, n = 108,788), morphology (22 traits, n = 294,533), and physiology (33 traits, n = 330,642).

The 375 phenotypic traits were further filtered for non-independence of traits, so that *p* values were merged for traits that were related to one another, resulting in a reduced data set of 226 traits, with a median sample size of 2,077 males and 2,435 females per trait.

### Linear mixed-effects models for static allometry

Our linear mixed-effects models indicated that 11 out of 226 traits (5%) (16 / 375 traits for unmerged p-values) are associated with scenario A (different slope, same intercept, Fig. 1A, 1D); most of these traits belonged to immunology and heart functional groups. Note that the intercept for each sex was set so that we compared mean values for each sex for a given trait. Scenario B (same slope, different intercept, Fig. 1B, 1E) was supported for 93 / 226 (41%) traits (165 / 375 traits for unmerged p-values). For scenario C (different slope, different intercept, Fig. 1C, 1F), 67 / 226 (30%) traits were categorized as consistent (81 / 375 traits for unmerged p-values), and the remaining 55 / 226 (24%) traits showed no significant differences in slope and intercept between males and females. Overall, when a statistically significant difference in allometric pattern was present between the sexes, intercept differences appeared more common than slope differences (41% compared to 5% traits), however both slope and intercept differences were also common (30%). Just under a fifth of traits showed no significant differences between males and females, indicating that, for most traits, sex differences in allometric patterning represent a significant source of variation in trait values.

**Figure 1.**
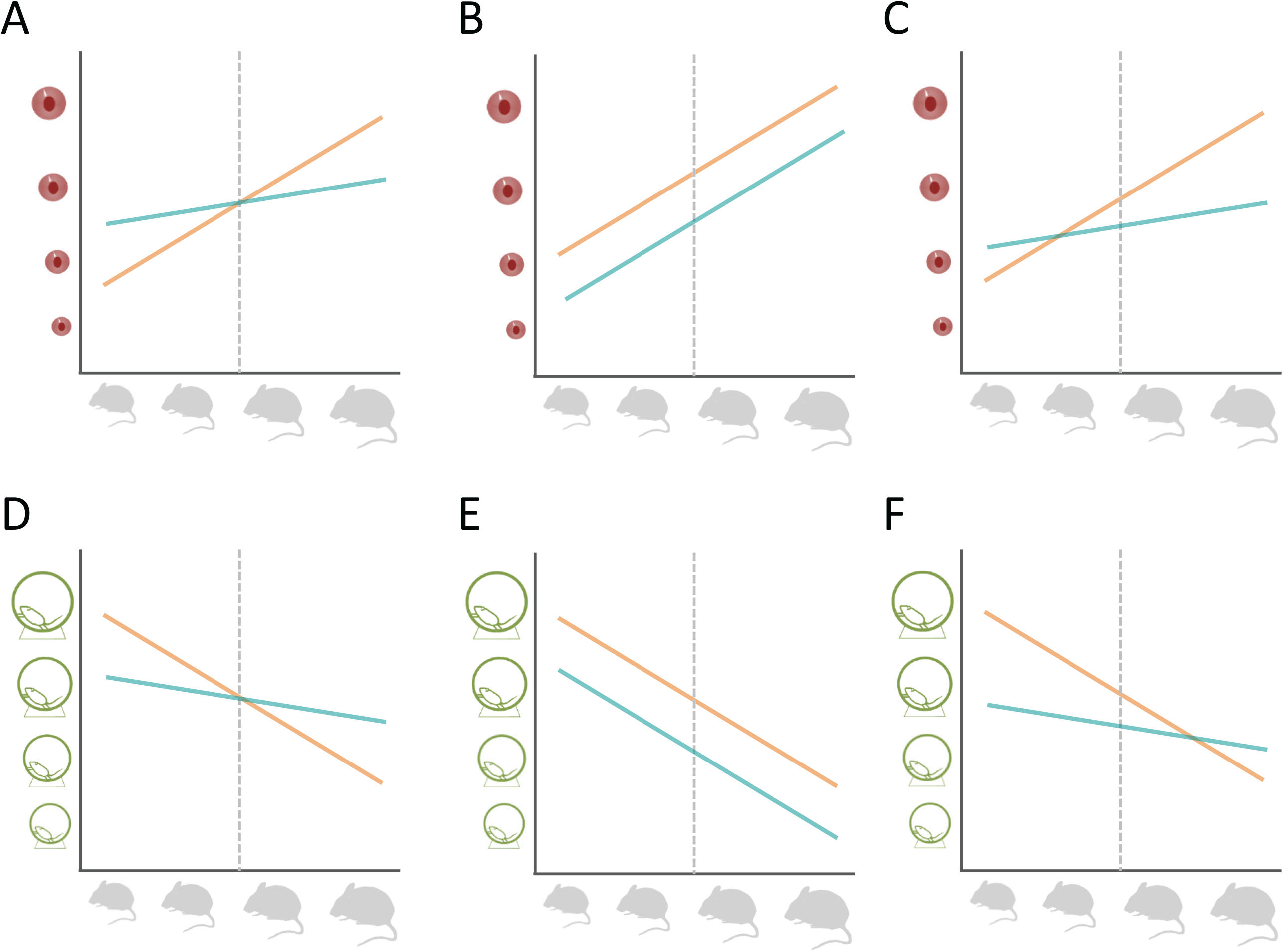
Examples of scenarios of sex differences in a trait of interest ∼ weight allometric relationship. Top row shows a hypothetical positive relationship between body weight and eye size and the bottom row negative relationship between body size and activity. Body weights are scaled and centred so that the intercept is at the trait mean represented by a grey dashed line. A) Different positive slopes for the sexes, but same intercepts. B) Same positive slopes for both sexes, but different intercepts. C) Different positive slopes for both sexes, and different intercepts. D) Different negative slopes for the sexes, but the same intercepts. E) Same negative slopes for both sexes, but different intercepts. F) Different negative slopes for both sexes and different intercepts.

Taken together, traits in all functional groups showed statistically significant (α = 0.05) sex differences. Slope differences between the sexes (scenario A) and intercept differences between the sexes (Scenario B) are most common in behaviour, immunology and physiology groups. Traits exhibiting both slope and intercept differences between the sexes (scenario C) were most commonly found in the behaviour, physiology and hematology functional groups. Non-significant differences in slope and intercept were most common among traits in the behaviour and eye functional groups.

### Sex bias in allometric parameters

Values for sex bias represent the number of traits that showed greater parameter value when males and females differed significantly. That is, we counted which sex displayed the greater intercept, slope and higher magnitude of variance. Sex bias in the slope and intercept values, in addition to the magnitude of variance (residual SD), showed considerable variability across functional groups, suggesting trait-specific patterning of sex differences. For scenario A, representing traits with significant differences in slope, most traits showed greater slope magnitudes for males (n = 10 traits), rather than for females (n = 6 traits) (Fig. 2A). For scenario B, females showed greater intercept magnitudes for heart, immunology, and eye functional groups (n = 65 traits), whereas males showed greater intercepts for traits in physiology, morphology, hematology, behaviour and hearing functional groups (n = 47 traits) (Fig. 2B). Overall sex bias (75 male traits: 90 female traits, Fig. 2B) was relatively lower for intercept differences, compared to slope differences (10 male traits: 6 female traits, Fig. 2A). Scenario C, which represents significant slope and intercept parameter differences between the sexes, was predominated by mixed bias across five out of nine functional groups (n = 29 traits), indicating that most functional groups contained traits that showed a mixture of directional differences in bias, comprising a combination of male bias in one parameter (slope or intercept) and female bias in the other parameter (slope or intercept) (Figure 2C). Eye-related traits represent an exception under scenario C, whereby traits with significant differences between the sexes did not show a mixed bias for slope and intercept values, consistent with few sex differences among traits. Across functional groups, male bias is slightly more common (5 groups) than female bias (4 groups) for statistically significant sex difference in residual SD, indicating that where traits show differences between the sexes, it is more common for males to be more variable than females, than *vice versa* (Figure 2D) (164 male traits: 97 female traits).

**Figure 2.**
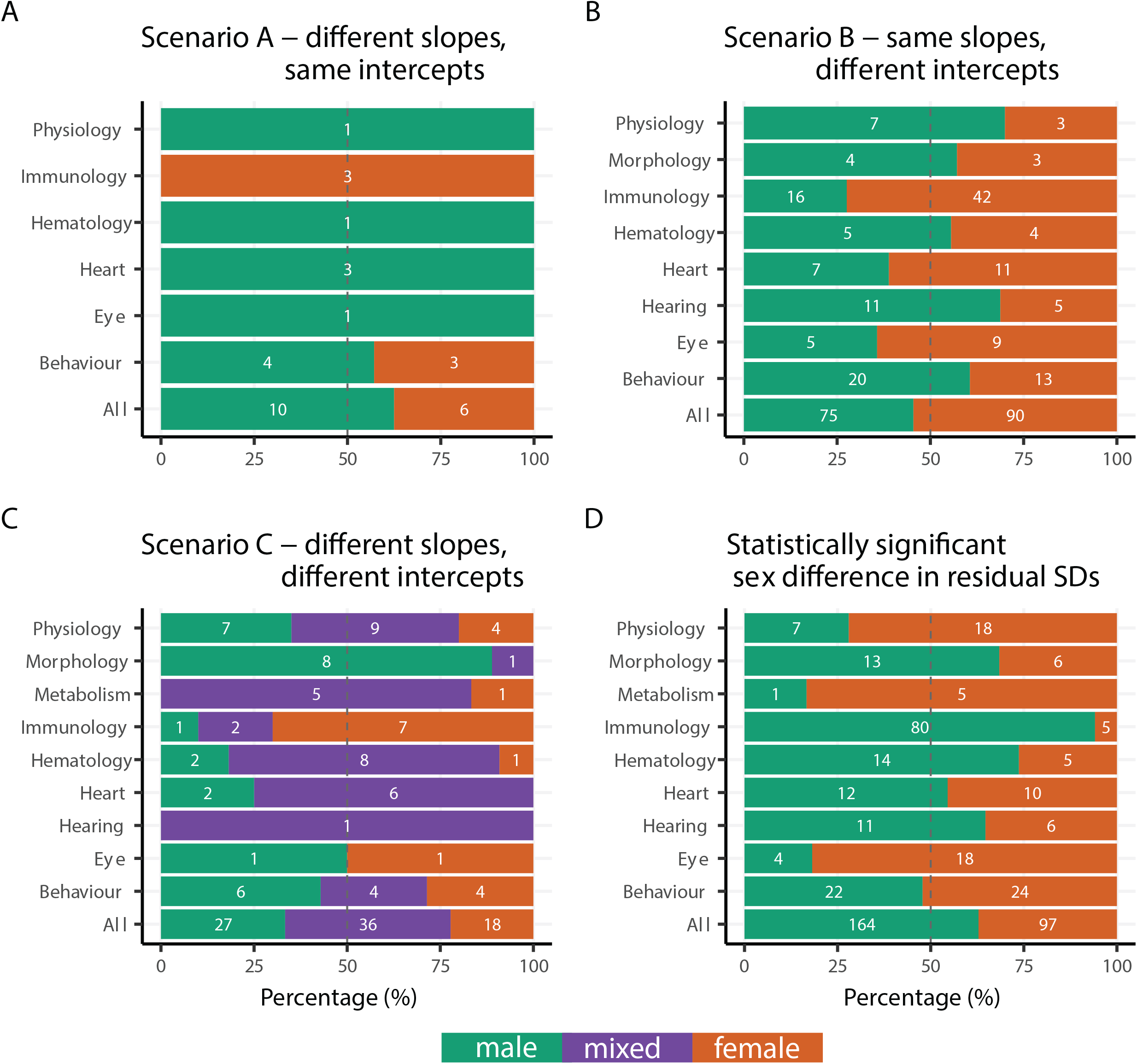
Sex biases for mice phenotypic traits arranged in functional groups. Sex bias represents a greater parameter value (slope, intercept, variance) in one sex compared to the other. Colours represent significant differences in trait values between the sexes (green – male biased, orange – female biased) for allometric slope (scenario A), intercept (scenario B) or slope and intercept, including traits with mixed (purple) significant differences (i.e. male-biased significant slope and female-biased significant intercept, or female-biased significant slope and male-biased significant intercept) (scenario C), and bias in statistically significant difference in variance (residual SD) between the sexes (D). The number of traits that are either female biased (relative length of orange bars) or male biased (relative length of green bars) are expressed as a percentage of the total number of traits in the corresponding group. Numbers inside the green bars represent the numbers of traits that show female bias within a given group of traits, values inside the orange bars represent the number of male biased traits, and those inside the purple bars represent a combination of female bias (for intercept or slope) and male bias (for intercept or slope).

### Meta-analysis and meta-regression of sex differences in slope, intercept, variance and model fit

Multi-level meta-analysis of absolute values in allometric slope and intercept, and variance revealed significant differences between the sexes (Fig. 3A – C), with the variation in the model fit (Fisher’s Zr^54^, transformed from marginal R^2^, variance accounted for by fixed effects) mimicking the variations in these three parameters of sex difference (Fig. 3D). Across functional groups, there was variability in the magnitude of absolute difference between the sexes, both within parameters (i.e., intercept) and across parameters. For absolute differences in intercept, traits within the metabolism functional group showed the greatest model point estimate difference between males and females, whereas those within the hearing group showed the smallest magnitude of difference (Fig. 3E). For differences in slope, which showed lower inter-trait variability than differences in intercept, the largest model point estimate difference was observed for immunology traits, and the smallest difference for hearing traits (Fig. 3F).

**Figure 3.**
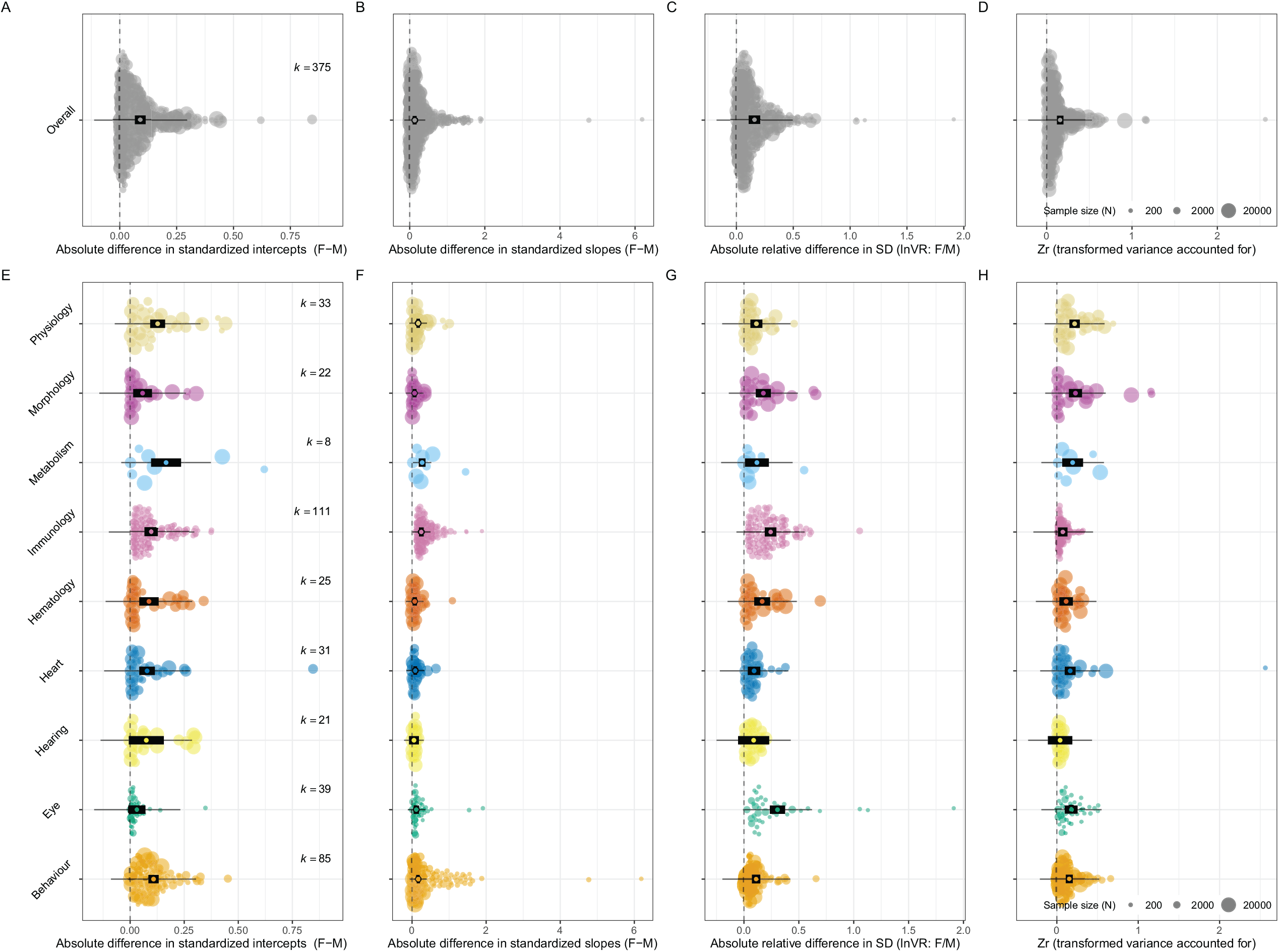
Orchard plots illustrating results of multivariate meta-analysis based on differences between male and female absolute values for allometric intercept (A, D), slope (B, E) and residual variance (SD) (C, F). Plots in greyscale (top row) show overall differences (A – C), and plots below, in colour, show separate results for each functional group (D – F). Orchard plots show model point estimate (black open ellipse) and associated confidence interval (CIs) (thick black horizontal line), 95% prediction intervals (PIs) (thin black horizontal line; PI represents heterogeneity), and individual effect sizes (filled circle), which are scaled by their sample size (N), the number of mice included per trait (see^53^). The number of effect sizes (number of phenotypic traits) is represented by *k*.

For the relative difference in residual SD, eye and immunology traits showed the largest amount of dimorphism, whereas hearing and metabolism traits were most similar in SD values between the sexes (Fig. 3G). Variance between the sexes accounted for by model fit (Zr) was highest in morphology and metabolism groups and lowest in the hearing group (Fig. 3H). Overall, across all parameters (intercept, slope, SD and model fit), confidence intervals (Cis) for hearing traits were the only ones to consistently overlap with zero, showing no statistically significant difference between the sexes (Fig. 3E-H). For traits within a given functional group, there was considerable variability in the magnitude of difference between the sexes. For sex differences in intercept, inter-trait variability was highest within physiology, metabolism and behaviour groups (Fig. 3E), whereas slope differences showed most inter-trait variability for eye and behaviour traits (Fig. 3F). Relative difference in SD was most variable among traits in eye and immunology groups (Fig. 3G), whereas sex differences in variance accounted for by model fit appeared most variable for heart and morphology groups (Fig. 3H).

### Relationship between slope/intercept, residual variance and model fit

Our quad -variate meta-regressions and ordinations of the relationships between slope, intercept and residual variance (Fig. 4) revealed weak, non-significant, correlations between either slope or intercept and residual variance (*r* = 0.04 – 0.09, Fig. 4A – B), indicating that a greater magnitude of difference between the sexes in either slope or intercept parameter is not strongly associated with greater trait variance. In contrast, absolute differences between the sexes in slope and intercept are strongly and significantly correlated (*r* = 0.74, Fig. 4C), indicating that in cases where there are significant differences in trait values for males and females, should a difference in intercept be present, this is likely accompanied by a difference in allometric slope.

**Figure 4.**
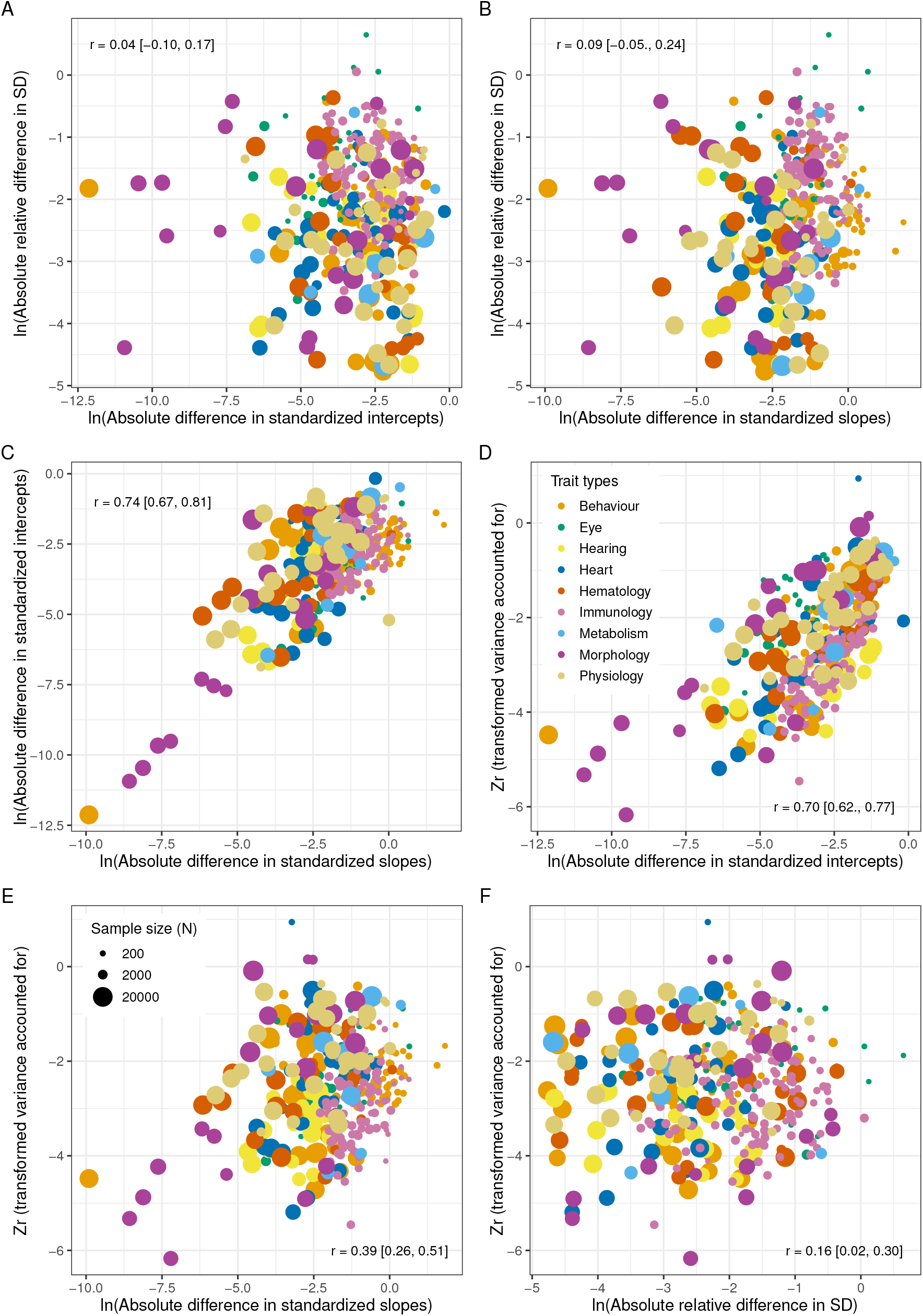
Bivariate ordinations of log absolute difference between males and females for intercept and residual SD (A), slope and residual SD (B), and slope and intercept (C) and between these three sex differences and the model fit for biological traits collated into nine functional groups (i.e., Trait types, represented as different circle colours). Individual effect sizes (circles) are scaled by their sample size (N), the number of mice included per trait.

Results from the quad-variate meta-regressions and ordinations of the relationships between slope, intercept and model fit (Zr) (Fig. 4D-F) indicated that both differences in intercept (*r* = 0.79) and slope between the sexes (*r* = 0.39) were significantly correlated with the model fit. As such, the greater the model fit (higher R^2^ marginal value, transformed to Zr) the lower the absolute difference between male and female values for intercept and, to a lesser extent, slope. In contrast, the difference in residual variance (Fig. 4F) was weakly, yet significantly, correlated (*r* = 0.16) with the model fit.

## Discussion

Most current medical guidelines are not sex-specific, being informed by preclinical studies that have been conducted only on male animals^4,10,55^ under the assumption that the results are equally applicable to females, or that the female phenotype represents a smaller body size version of the male phenotype^21,56^. We find that 76% of traits are dimorphic, either in slope, intercept or both slope and intercept, and that among these 41% of traits show scaling differences between males and females. Scaling differences are not supported across all phenotypic traits, indicating that body weight alone is not enough to explain sex differences in phenotypic traits but that it may be useful for some traits.

In an era where personalised medicine interventions are within reach and patient-specific solutions represent a realisable frontier in healthcare (e.g.,^57-59^) it is now well recognised that sex-based data are much needed to advance care in an equitable and effective manner. The historic neglect of sex as a study variable means that the natural history and trajectory of treatment response in women remains opaque for many chronic diseases. As studies that illuminate the presence and importance of sex differences continue to emerge, many experimental set-ups that use both sexes continue to eschew downstream testing for sex differences, in part due to perceived inflation of sample size required for such analyses^56,60-62^.

Explicit male-female comparisons are needed to clarify the nature of sex differences^55,63^. Here we address this issue through a novel meta-analytical focus on identifying and characterising allometric scaling relationships for biological traits on a broad scale. We identify slope parameter (*b*) differences between the sexes, and find these are mostly associated with significant differences in intercept value (Fig. 4C). We therefore demonstrate that the relationship between trait and body mass in mice differs fundamentally in mode (i.e., change in inter-trait covariance) between the sexes and that dimorphism cannot be fully explained by a magnitude shift in intercept value, as would be predicted should female phenotype represent a scaled version of male phenotype. For traits where there are significant differences in both slope and intercept between the sexes, it is common for a mixed scenario (male-biased significant slope and female-biased significant intercept, or female-biased significant slope and male-biased significant intercept; note that intercepts represent mean values for each sex) to occur. At least 35% of traits show differences not accounted for by scaling alone, meaning a female value cannot be predicted based on an allometric coefficient extracted from regression data collected on males for over a third of traits. Further, we find a male bias in residual SD for traits in morphology, immunology, hematology, heart, and hearing functional groups (7 out of 9 functional groups), indicating greater variance in males compared to females for these traits.

Our results complement recent evidence that supports a complex, trait-specific patterning of sex differences in markers routinely recorded in animal research^18,20,64^. Specifically, we build on previous studies using phenotypic traits from the International Mouse Phenotyping Consortium that have identified that sexual dimorphism is prevalent among phenotyping parameters^20^, and moreover that, contrary to long-held assumption, neither females nor males show greater trait variability. We here show that the allometric relationship between trait value and body weight is dimorphic for most traits (76%), and these differences, where present, reflect trait-specific allometric patterns, involving both slope and intercept changes. As such, for slopes greater than zero, some trait values increase faster than body weight (positive allometry; *b* > 1) and some do not increase at the same rate as body weight (negative allometry; *b* < 1).

### Sex-based scaling in biomedical studies

We asked the question of whether all or some sex differences in phenotypic traits are due to differences in body weight. Our finding, that some but not all traits scale between males and females (41%, Fig. 2B, 3E), has implications for drug therapy, and specifically data surrounding the efficacy of drug dosing scaled by body weight.

There exist known sex differences in drug prescription prevalence and usage patterns, as well as response to drug therapy^65,66^. The same therapeutic regimen can elicit different responses due to sex-specific variance in pharmacokinetics and pharmacodynamics profiles (e.g.,^67,68^), arising from underlying physiologic differences. These include, for example, significantly dimorphic traits captured among the physiology group in our analysis, such as iron^69^ and body temperature^70^, among the morphology group, such as lean mass and fat mass^65^, and among the heart functional group, such as QT interval (time between Q wave and T wave)^71^. Further, women are 50 – 75% more likely to experience Adverse Drug Reactions (ADRs)^73^, although these are not fully explained^24^. Women may be at increased risk of ADRs because they are prescribed more drugs than men, however women are usually prescribed drugs at the same dose as men, meaning that they receive a higher dose relative to body weight in most cases. Scaling of doses on a milligram/kilogram body weight basis has been recommended as a pathway to reducing ADRs^23^, particularly for drugs that exhibit a steep dose-response curve^74^. Indeed, sex differences in ADRs have been argued to be the result of body weight rather than sex, per se^22^. For both assertions to be supported, we would expect to observe a scenario (here, scenario B) whereby most or all phenotypic traits exhibit a scaled relationship between males and females, as a function of body weight.

Our results indicate that 41% traits follow scenario B, with many traits (35%) also supporting a sex- and trait-specific relationship between weight and phenotypic trait. This aligns with evidence that weight-corrected pharmacokinetics are not directly comparable in men and women^23,75^, and that many sex differences in ADRs persist after body weight correction^76,77^. Nevertheless, the Food and Drug Administration (FDA) has recommended dosage changes for women (e.g., sleep drug zolpidem^78^) and weight adjusted dosing of some drugs, such as antifungal drugs and antihypertensive drugs, which appear to ameliorate sex differences in pharmacokinetics^79,80^. Our results are consistent with support for scaling in some circumstances, as we find 41% traits follow this relationship. The greatest number of these traits occur in the immunology and heart groups, which contain parameters most closely relevant to pharmacokinetic and pharmacodynamic factors, compared to the behaviour group, where many scaling differences are found but translate less closely to human behaviour relevant to therapeutic drug use. We suggest traits in scenario B as candidates for further investigation in weight-adjusted dosing. If weight-adjusted dosing is supported based on intercept scaling (scenario B), the extent to which variation in the allometric relationship may reduce the efficacy of such an approach also requires consideration. This is measured by model fit in our analyses, whereby greater model fit equates to more of the variance in phenotypic traits for males and females being explained by body weight. For those traits that scale (scenario B), the most useful candidates for dose-scaling are therefore likely to be captured by those traits with the highest model fit (Zr, transformed from R^2^ marginal) (Fig.4D). Traits with lower model fit may scale, but a considerable amount of variance in the phenotypic trait is not explained by body weight, therefore scaling may not achieve the desired clinical outcome. Traits within scenario B that meet this criterion are most frequently found in the eye, morphology and physiology groups.

We suggest that where there exists an association between sex and dose, dose-response curves are likely to be sex-specific and clarification of this relationship would be supported (e.g., using meta-analysis^81^). Since many drugs are withdrawn from the market due to risks of ADRs in women, meta-analytic approaches to illuminating sex-specific dose response curves represents a viable opportunity to reducing the number of ADRs and reaching an important target set by precision medicine^82^. We further note that human behaviour, an important variable that impacts drug efficacy and ADRs, is not well-represented by our mouse data. Behavioural factors such as differences in health-seeking behaviours and prescription patterns may relate to a higher prevalence of use for most therapeutic drugs in women as compared to men^66,72^.

### Implications for allometric evolution

The study of allometry has a long history in evolutionary biology, established as a foundational descriptor of morphological variation at ontogenetic, population and evolutionary levels^83,84^. Allometry may channel phenotypic variation in fixed directions, defining scaling relationships that persist across large evolutionary timescales. For example, craniofacial variation among mammals has been observed to be constrained by allometry, such that small mammals have shorter faces than do larger ones^85^. Conversely, allometry may facilitate morphological diversification, acting as a line of ‘evolutionary least resistance’, allowing for new morphotypes to originate relatively rapidly among closely related species^48,86^. These pathways (allometric constraint vs allometric facilitation) may be a start point for exploring how sex differences in disease phenotypes arise, data that have been cited as a potential unexploited resource relevant for the development of new therapies^87^.

Studies of static allometry, as examined herein, have revealed low levels of intraspecific variation in allometric slope, which explains only a small proportion of variation in size^88^, compared to variation in allometric intercept^89^. Moreover, traits under sexual selection have also revealed low magnitudes of allometric slope change under artificial selection experiments^90^ and in wild populations^91^, whereas intercept changes appear clear and heritable. These differences have historically been thought to be due to underlying features of the developmental system acting as an internal constraint^46,92^, whereas more recent interpretations suggest that external constraint (selection) more likely acts to maintain slope invariance at the static level^48^, which is consistent with data showing that variation occurs instead at the ontogenetic level, i.e. growth rate and ontogenetic allometric slope are evolvable (e.g.,^93-95^). Broadly consistent with other static allometric studies, we find that where differences in allometry are present, significant intercept shifts alone are more common than are significant slope shifts (Fig. 2A compared to 2B). We focus explicitly on sex differences and observe that many traits show a combination of intercept and slope changes, as well as differences in residual variance. Aside from the evolutionary implications – that allometric slope likely does not have a high evolvability, or capacity to evolve – many of the traits examined here may show a low level of sex difference in slope because the sexes are both experiencing the same selective pressure to maintain functional size relationships across different body sizes.

Our meta-analytic results build a narrative of complexity in sex-based trait interactions and promote a case-specific approach to preclinical research that seeks to inform drug discovery, development and dosage. Across a diverse set of phenotypic traits we show that differences in body weight are not sufficient to explain sex differences in trait values, but scaling differences are common and body-weight scaling may be helpful in some traits. Our results evidence the plasticity of allometry at a microevolutionary scale, revealing a pathway for sex variation in phenotypic traits, which may influence study outcomes in biomedicine.

## Methods

### Data compilation and filtering

We conducted all data procedures, along with statistical analyses, in the R environment v. 4.1.396. We compiled our data set from the International Mouse Phenotyping Consortium (IMPC) (www.mousephenotype.org, IMPC data release 10.1 June 2019), accessed in October 2019. These represent traits recorded in a high-throughput phenotyping setting whereby standard operating procedures (SOPs) are implemented in a pipeline concept. The phenotypic traits represent biomarkers used for the study of disease phenotypes (see^20^), collated into the following nine functional groups: behaviour, eye, hearing, heart, hematology, immunology, metabolism, morphology, and physiology, which are the IMPC’s original categorization (also previously used in Zajitschek et al.^18^). These groupings were assigned in relation to the description of the procedure undertaken for data point collection and following the categorisation of pipeline events at adult stage, detailed in the International Mouse Phenotyping Resource of Standardised Screens (IMPReSS, https://www.mousephenotype.org/impress/index).

For the initial dataset, data points were collated for adult wildtype mice only, filtering to include non-categorical phenotypic trait values for which covariate information on sex and body weight were available. Note that all mice were from the C57BL/6 strain, but they come from seven sub-strains. This initial dataset comprised of 2,866,345 data points for 419 traits. A series of data cleaning procedures were implemented to remove data points with missing body weight, zero values for a phenotypic trait and duplicated specimen IDs. Data filtering was conducted using the R package dplyr v.1.0.7^97^. The resulting data set comprised 2,118,370 data points for 379 phenotypic traits, all of which had corresponding body weight data, enabling us to estimate an allometric relationship between a trait of interest and body weight. Of these, 89 traits (24%) were on the interval scale, and were therefore adjusted to be on the ratio scale. For each phenotypic trait, we had the following variables (covariates): phenotyping center name (location where experimental data were collected), external sample ID (animal ID), metadata group (identifier for experimental conditions in place during the experiment), sex (male / female), weight (body weight in grams), weight days old (day on which weight was recorded), strain name (identifier character for mouse sub-strains), procedure name (description of the experimental procedure as in IMPReSS), parameter name (description of the recorded parameter as in IMPReSS), and data point (phenotypic trait measurement – response variable).

### Linear mixed-effects model for static allometry

The static form of allometry, the covariation of a trait with size as measured across a population of adults within a single species^84^, was quantified using a linear mixed-effects model approach^98^. Within this framework, the relationship between phenotypic trait value and body weight, accounting for random effects associated with assignment to a metadata group and batch (defined as the date when the measurements are collected), was quantified for each of the 297 traits. Models were constructed using the function *lme* in the R package nlme v. 3.1-153^99^ and applied to each phenotypic trait separately. We used the approach described by Nakagawa et al.^100^ that uses within-group centring (wgc) of the continuous predictor (i.e., weight); in this way, the intercepts (*x* = 0) for each sex represents the population mean for that specific sex. Also, we calculated z-scores (*z*) from the response (y) so that all regression coefficients are directly comparable across different traits. The applied model was:

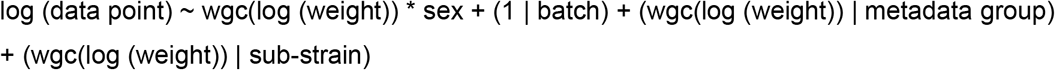

The random factor ‘batch’ labelled a cohort of mice that went through a procedure on the same day (see^20^), ‘metadata group’ represented occasions when procedural parameters were changed (e.g., different instruments, different observers and different settings) and strain represents the mouse strain identifier (e.g., C57BL/6N). These three random factors along with the ‘weight’ random slopes would reduce Type I errors due to clustering^101^. Also, to estimate different residual variances between the two sexes, we modelled group-wise heteroscedasticity structure, which was defined using the *lme* function’s argument weights = varIdent (form = ∼1 | sex). We parsed a custom function to apply this model using conditional statements to account for situations where the random factor (sub-strain or metadata group) has only one unique term (e.g., data for trait ‘tibia length’ comprised only of strain C57BL/6N). In other words, some traits had only one sub-strain or one meta-data group, and in such instances we did not fit the corresponding random effect. This procedure resulted in a final data set of 2,105,419 and 375 traits (i.e., the model did not converge for four traits, which were excluded from the subsequent analysis).

For each phenotypic trait, model parameters (regression coefficients and variance components) were extracted, using R package broom.mixed v.0.2.7^102^, for males and females (slope, intercept, standard error, SE of slope, SE of intercept, R^2^ marginal, R^2^ conditional and residual variance) and corresponding *p* values for regression coefficients were extracted to assess the significance of sex differences in slope and intercept. Because the *lme* function did not provide statistical significance for differences in residual variances (standard deviations, SDs), we used the method developed by Nakagawa et al^103^ or the logarithm of variability ratio, which compares the difference in SDs between two groups to obtain *p* values for residual SD differences (see also Senior et al.^104^).

We were aware that some of the 375 studied traits were strongly correlated (i.e., non-independent: e.g., traits from left and right eyes and immunological assays with hierarchically clustering and overlapping cell types). Therefore, we collapsed *p* values of these related traits into 226 *p* values, using the procedure (grouping related traits or trait grouping) performed by Zajitschek and colleagues^18^. We employed Fisher’s method with the adjustment proposed by Li and Ji (2005) implemented in the R package, poolr^105^, which modelled the correlation between traits; we set this correlation to 0.8.

### Static allometry hypotheses and Sex-bias in allometric parameters

Using parameters extracted from the above models, three scenarios were assessed (see Fig 1), describing the form of sex differences in the static allometric relationship between phenotypic trait value and body weight. For a given trait, these were: a) males and females have significantly different slopes but share a similar intercept (Fig. 1A, 1D), b) males and females have significantly different intercepts but share a similar slope (Fig. 1B, 1E), c) males and females have significantly different slopes and intercepts (Fig. 1C, 1F). In addition, we assessed how many traits were significantly different in residuals SDs between the sexes. For these classifications, we used both *p* values from 375 traits and 226 merged trait groups.

For scenarios A – C, which represent significant differences between male and female regression slope and / or intercept parameters and cases where sex differences in SDs were significant, data were collated into functional groupings (as listed above) to assess whether, and to what extent, sex bias in parameter values and variance was present across phenotypic trait values. That is, when males and females differed significantly, we counted which sex displayed the greater parameter value (intercept, slope) and, separately, we also tallied the sex with the higher magnitude of variance. Results were pooled for phenotypic traits within a functional group and visualised using R package ggplot 2 v. 3.3.5^106^ for scenarios A – C, resulting in one set of comparisons for parameter values, and one for variance (SD) values. We should highlight that we only used the data set with 375 traits because the directionality of some trait values became meaningless once traits were merged, although merged *p* values were meaningful as *p* values are not directional (e.g., spending time in light side or dark side).

### Meta-analysis of differences in slopes, intercepts, residual SDs and model fits

We were aware that our classification approach using *p* values is akin to vote counting, which has limitations^107^. Therefore, we conducted formal meta-analyses using the following effect sizes: 1) difference between intercepts (traits mean for males and females), 2) difference between slopes, and 3) differences between residuals SDs used corresponding SE or, more precisely, the square of SE as sampling variance. We were not able to compare the directionality of effect sizes among traits (e.g., latencies and body sizes), however our main interest in this study was whether males and females were different in intercepts, slopes and residuals SDs irrespective of directionalities. Therefore, we conducted meta-analyses of magnitudes applying the transformation to the mean and sampling variance, which assumes to follow folded normal distribution (eq. 8^108^), by using the formulas below:

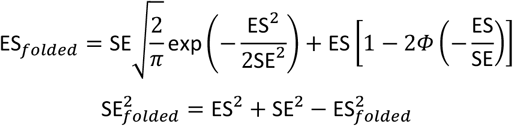

Where Φ is the standard normal cumulative distribution function and ES*folded* and SE*folded* are, respectively, transformed effect size (point estimate) and sampling variance, while ES and SE are corresponding point estimate and sampling variance before transformation. In addition to these three effect sizes, we also meta-analysed another effect size, which is the model fit quantified by marginal R^2^ (variance accounted for by fixed effects; in our case, weight, sex and their interaction). The marginal R^2^ values were squared-root transformed into correlation values, which are akin to the corelation between observed trait values and model predicted values (based on fixed effects). Then, we transformed this correlation value to Fisher’s Zr so that we could calculate sampling variance based on sample sizes.

Morrissey^108^ has shown that meta-analytic means using such a folding transformation (absolute valued effect sizes) are hardly biased. Therefore, these transformed variables were directly meta-analysed using the *rma*.*mv* function in the R package, metafor^109^. The intercept models (meta-analytic model) had three random factors: 1) functional group, 2) traits group and 3) effect size identifier (which is equivalent to residuals in a meta-analytic model^110^), while in the meta-regression models, we fitted functional group as a moderator (see Fig 3). The model structures for all the three effect sizes were identical. We reported parameter estimates and 95% confidence intervals, CI and 95% prediction intervals, PI, which were visualised by the R package, orchaRd^53^. In a meta-analysis, 95% PI represents the degree of heterogeneity as well as a likely range of an effect size for a future study. We considered the estimate statistically significant when 95% CI did not span zero.

### Correlations among differences in slopes, intercepts, residual SDs and model fits

We also quantified correlations among the four effect sizes, using a Bayesian quad-variate meta-analytic model, implemented in the R package, brms^111^. We fitted functional grouping as a fixed effect and trait groups as a random effect using the function, *brm*. Notably, we have log transformed ES*folded* and Zr and also transformed SE*folded* and SE for Zr using the delta method (e.g.,^112^), accordingly, before fitting effect sizes to the model. We imposed the default priors for all the parameter estimated with the settings of two chains, 1,000 warm-ups and 4,000 iterations. We assessed the convergence of the chains by Gelman-Rubin statistic^113^, which was 1 for all chains (i.e., meaning they were all converged) and we also checked all effective sample sizes for posterior samples (all were over 800). We reported mean estimates (correlations among the three effect sizes and model fits) and 95% credible intervals (CI) and if the 95% CI did not overlap with 0, we considered the parameter statistically significantly different from 0.

## Data availability

Source data are provided with this paper. The R code and data generated during this study are freely accessible on GitHub at [to be updated on acceptance].

## Code availability

An R Markdown file with instructions and the complete workflow for all analyses is provided in the supporting information, available at [to be updated on acceptance].

### Acknowledgements

This research was supported by Australian Research Council grants DP200100361 awarded to SN and ML and FT200100822 awarded to LABW. Research reported in this publication was supported by the European Molecular Biology Laboratory core funding and the National Human Genome Research Institute of the National Institutes of Health under Award Number UM1HG006370. The content is solely the responsibility of the authors and does not necessarily represent the official views of the National Institutes of Health.

## Author contributions

LABW and SN designed the research; SN, LABW, SRKZ, ML and HH contributed to the conception and implementation of data analysis; JM contributed to data acquisition; LABW drafted the manuscript with contributions from SN and ML.

